# High Resolution Spatial Mapping of Microbiome-Host Interactions via *in situ* Polyadenylation and Spatial RNA Sequencing

**DOI:** 10.1101/2024.11.18.624127

**Authors:** Ioannis Ntekas, Lena Takayasu, David W. McKellar, Benjamin M. Grodner, Chase Holdener, Peter Schweitzer, Maya Sauthoff, Qiaojuan Shi, Ilana L. Brito, Iwijn De Vlaminck

**Author notes:** These authors contributed equally (Ioannis Ntekas, Lena Takayasu, David W. McKellar).

## Abstract

Inter-microbial and host–microbial interactions are thought to be critical for the functioning of the gut microbiome, but few tools are available to measure these interactions. Here, we report a method for unbiased spatial sampling of microbiome-host interactions in the gut at one micron resolution. This method combines enzymatic *in situ* polyadenylation of both bacterial and host RNA with spatial RNA-sequencing. Application of this method in a mouse model of intestinal neoplasia revealed the biogeography of the mouse gut microbiome as function of location in the intestine, frequent strong inter-microbial interactions at short length scales, shaping of local microbiome niches by the host, and tumor-associated changes in the architecture of the host-microbiome interface. This method is compatible with broadly available commercial platforms for spatial RNA-sequencing, and can therefore be readily adopted to broadly study the role of short-range, bidirectional host-microbe interactions in microbiome health and disease.

## INTRODUCTION

It has long been speculated that the gut microbiome functions as an organ system with tissue-like properties defined by dynamic interactions between microbial and host cells^1,2^. Yet, investigating the tissue-properties of the gut microbiome has been difficult due to a lack of adequate measurement tools^3^. While advances in imaging have enabled the study of the localization of specific microbes in the gut^4^, these methods are limited in multiplexity or fail to provide detailed information about host function and response^3–7^. Spatially resolved RNA-sequencing (RNA-seq), a more recent approach to studying gene expression in tissues, has been used to examine the cellular architecture of intestinal tissues in health and disease^8–12^. Nevertheless, characterizing the microbiome-host interface via spatial RNA-seq remains challenging due to constraints in spatial resolution and sensitivity to microbial RNA^13^. Moreover, existing approaches rely on the spurious capture of A-rich microbial RNAs via poly(dT) primers or use a limited set of microbe-specific primers, which leads to measurement biases and a limited scope of discovery^13–16.^

Here, we address these limitations by exploring the use of enzymatic polyadenylation of microbial RNA and host RNA *in situ* to map the microbiome-host interface via spatial RNA-seq (**Fig. 1**)^17^. We demonstrate that enzymatic *in situ* polyadenylation significantly improves bacterial RNA recovery by oligo(dT) based spatial transcriptomics arrays, by up to 100-fold, and we show that this chemistry is compatible with multiple commercially available platforms for spatial RNA-seq. The enhanced recovery of bacterial RNAs enables dense spatial sampling of the microbiome at single micron resolution<u>^1^</u>. In addition to bacterial RNAs, *in situ* polyadenylation enables capture and characterization of both A-tailed and non-A-tailed transcriptomes of host cells within the intestine. By integrating these layers of information, the resulting spatial RNA-seq method provides a highly detailed view of microbiome-host interactions in the gut (**Fig. 1**). Application of this method revealed the location-dependence of the organization of the microbiome in the mouse intestine, interactions within and between microbial taxa at short length scales, local shaping of the microbiome by the host via immune and antimicrobial signaling, and changes in microbiome and host cell architectures at microbiome-tumor interfaces.

**Figure 1.**
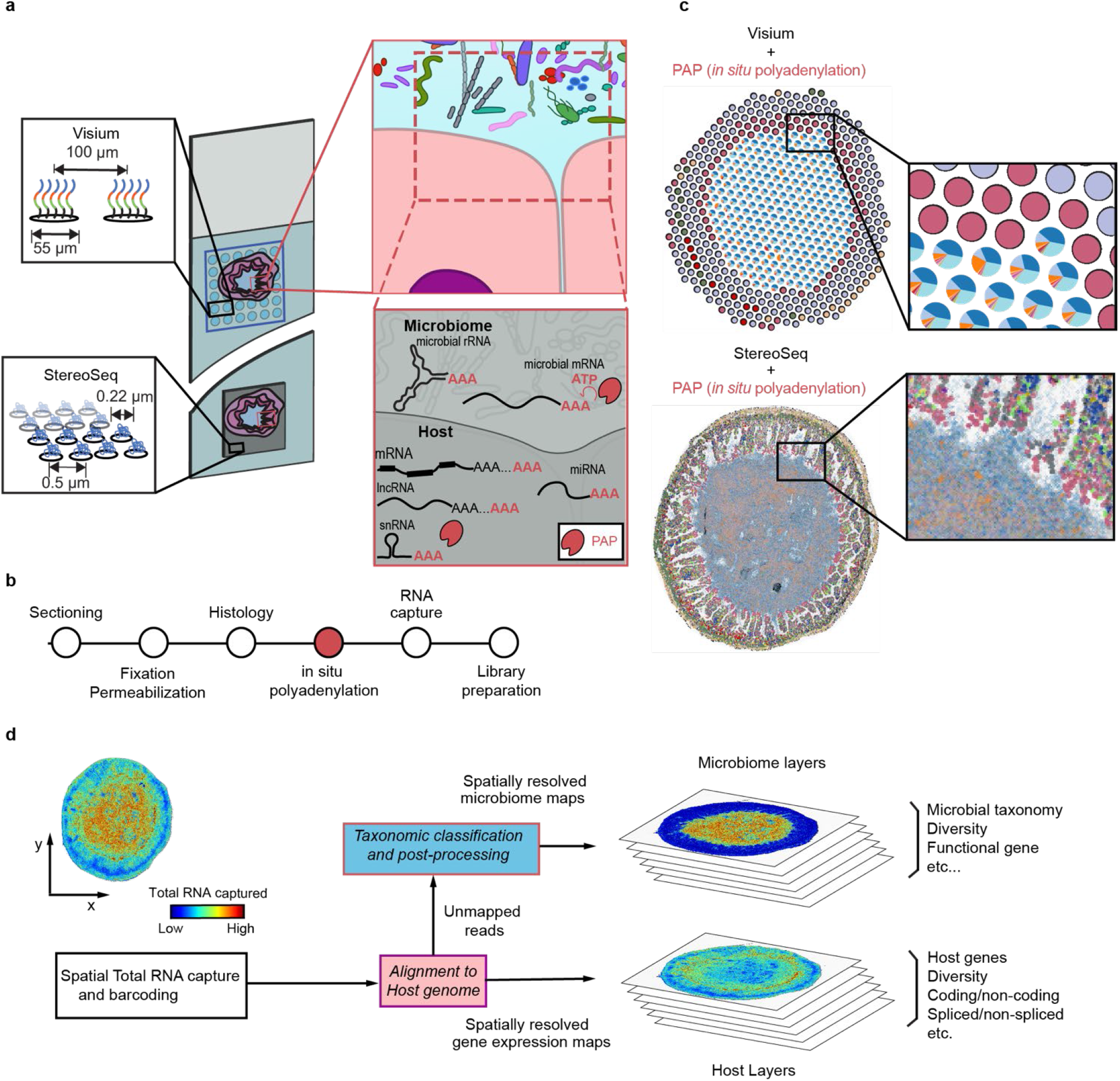
*In situ* polyadenylation enables the capture of microbiome signals with sequencing-based spatial transcriptomics platforms. **a**. Overview of the experimental design. Array-based spatial RNA sequencing (at low or high spatial resolution) is combined with *in situ* polyadenylation via Poly(A) polymerase (PAP). **b**. Schematic of protocol for microbiome and host Spatial Total RNA-Sequencing. The standard steps of cryosectioning, fixation, and histology are followed by enzymatic *in situ* enzymatic polyadenylation, total RNA capture, and sequencing library preparation. **c**. Example data for the low (top) and high (bottom) resolution platforms. **d**. Schematic of bioinformatics workflow.

## RESULTS

### Spatial mapping of microbiome-host interaction via in situ polyadenylation

We tested whether *in situ* polyadenylation is an effective method to enhance the recovery of microbiome-derived RNA and map host-microbiome interactions, initially at low spatial resolution. We collected fresh-frozen tissue from a mouse model of colorectal cancer (APC-deficient) at four distinct locations: proximal small intestine, ileum, cecum, and colon, and performed spatial transcriptomics on the Visium platform (**Fig. 2a**). Immediately after sectioning, we fixed the tissues with methacarn, which we found to be important for retaining fecal content in the gut sections (**Methods**). Following fixation, H&E staining, and imaging, we performed *in situ* enzymatic polyadenylation to enable the capture of non-polyadenylated molecules, including non-coding RNA and microbial RNA. To quantify the effect of enzymatic polyadenylation, we also performed conventional spatial RNA-seq, without *in situ* polyadenylation, on proximal tissue sections (**Methods**). After cDNA synthesis and sequencing, we obtained an average of 156 million reads per sample (156 M ± 43 M).

**Figure 2.**
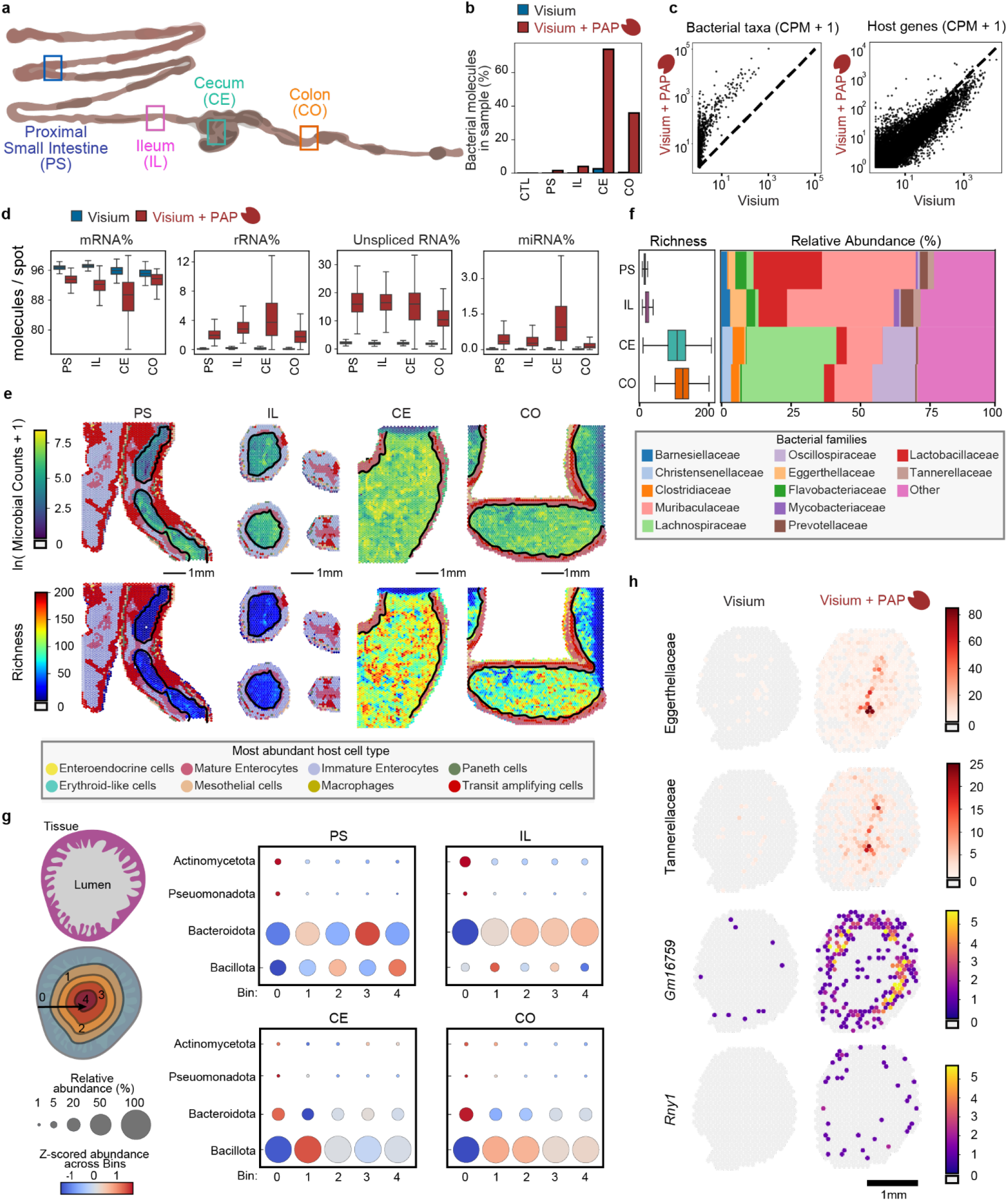
Spatial Total RNA-Sequencing of the murine Gastrointestinal (GI) tract with the Visium platform. **a**. Sampling locations across the murine GI tract measured with and without *in situ* polyadenylation (Proximal small intestine (PS), Ileum (IL), Cecum (CE), and Colon (CO)). **b**. Barplots showing the percent of Unique molecules classified as bacterial in the paired experiments with and without *in situ* polyadenylation. CTL = murine heart tissue

To quantify microbial and host-specific sequences, we initially mapped the reads to the murine (host) reference genome and then performed taxonomic classification on the unmapped reads using Kraken2^18^ (**Methods**). To assess potential contamination and sequence misclassification, we analyzed non-intestinal tissue (murine heart). We found very low levels of microbial signal in these non-intestinal tissues (0.002-0.04 % of total reads classified as microbial, with and without the polyadenylation step, **Fig. S1**). We next quantified the enrichment in microbial RNA enabled by *in situ* polyadenylation. We found that *in situ* polyadenylation resulted in up to a 99-fold enrichment of bacterial RNA (**Fig. 2b**), with improved capture for most microbial taxa, while maintaining high capture efficiency for host genes (**Fig. 2c, Fig. S2a**). The enrichment of RNA from viruses and archaea was greatest in the proximal small intestine (10-fold and 6-fold increase, respectively, **Fig S2b-c**). Notably, *in situ* polyadenylation enhanced detection of both lowly abundant bacterial taxa (e.g. Tannerellaceae and Eggerthellaceae families) and highly abundant taxa (e.g. Lactobacillaceae and Lachnospiraceae). In contrast, conventional spatial RNA-seq (Visium) captured a limited diversity and often failed to detect microbial RNA even in the center of the lumen **(Fig. S3**).

In addition to microbial RNA, we found that *in situ* polyadenylation also improved the capture of host-derived non-polyadenylated RNAs (**Fig. 2d and Fig S4**). For example, unspliced mRNAs were enriched after polyadenylation (15.6% of unique molecules vs 2.1%, **Fig. 2d**). These unspliced molecules likely represent nascent transcripts, which could provide insights into cellular responses to microbial cues and into cellular turnover and replenishment. Other biotypes were enriched, including ribosomal RNAs (rRNAs, 2.6% versus 0.16%), microRNAs (miRNAs, 0.622% vs 0.021%), small nucleolar RNAs (snoRNAs, .0683% vs 0.009%), long non-coding RNAs (lncRNAs 2.80% vs 1.42%), small nuclear RNAs (snRNAs 0.0425% vs 0.0012%), and miscellaneous RNAs (miscRNAs, 0.2557% vs. 0.001%). *In situ* polyadenylation enabled the identification of RNAs that are common to all four GI tract regions and murine heart tissue, including miscRNAs such as *Rny1* and *Rny3*, the vault RNA *Vaulrc5*, and the snRNA *Rn7sk* (**Fig S4c**). Additionally, we observed molecules with spatially patterned expression, including the lncRNA *Gm16759*, which was enriched specifically in the ileum. *Gm16759* has been shown to regulate *Smad3* expression, inhibiting the induction of intestinal regulatory T cells via the TGF-β pathway^19^. In the proximal intestine, we observed expression of the lncRNA *Gm31992*, while in the distal sections, including the cecum and large intestine, we detected expression of other non-coding features including the lncRNAs *Gm56583* and *miR9-3hg*, which is implicated in human cancer^20,21^.

We next examined microbiome composition as a function of location within the gastrointestinal (GI) tract. Moving down the GI tract from the proximal small intestine and ileum to the cecum and colon, we observed an increase in taxonomic richness per spot (average of 14.1 genera in the small intestine to 114.4 in the large intestine, lumen) (**Fig. 2e**). Lactobacillaceae and Muribaculaceae were abundant in the proximal small intestine (PS) and ileum (IL), but not in the cecum (CE) and colon (CO). Lachnospiraceae and Clostridiaceae had the greatest abundance in the cecum while Oscillospiraceae had the highest abundance in the colon. Last, Flavobacteriaceae, Eggerthellaceae, Barnesiellaceae, Prevotellaceae, and Tannerellaceae had higher relative abundances in the small intestine (**Fig. 2f**). These results are in line with the previous findings^22^.

A major advance of the method is its ability to examine changes not only along the longitudinal axis but also along the transverse axis of the GI tract, from the tissue to the lumen, where variations in micro-niches—such as pH, oxygen levels, nutrient accessibility, and contact with the host’s defense mechanisms—are expected to influence microbial composition^22^. In the small intestine, we observed that the microbial signal originates near the center of the lumen, where it also becomes more diverse. In contrast, in the cecum and large intestine, we observed a strong microbial signal and increased diversity near the mucosa. The limited resolution of the Visium platform used in this experiment, however, did not permit to fully resolve the mucosal layer or the interface between the lumen and mucosa (**Fig. 2e**). To further assess changes in microbiome composition along the transverse axis, we divided each map into five bins based on distance to the lumen. We then measured the relative abundance in each bin for four representative phyla: Actinomycetota, Pseudomonadota, Bacteroidota, and Bacillota (**Fig. 2g**). Doing so, we found that Actinomycetota and Pseudomonadota were generally more abundant near the mucosa and tissue layer, particularly in the small intestine. Bacteroidota, the most abundant phyla in the small intestine, were preferentially present away from the tissue and mucosa. Conversely, Bacillota were the dominant phyla in the cecum and large intestine across all bins, with higher levels observed away from the tissue, while Bacteroidota were enriched in the tissue layer (**Fig S3b**). These results demonstrate the effectiveness of *in situ* polyadenylation for spatially mapping microbiome-host interactions, enriching the capture of non-host and non-coding molecules (**Fig. 2h**), and inspire experimentation at higher spatial resolution.

### Mapping microbiome-host interaction at higher spatial resolution

We next implemented *in situ* polyadenylation on a high-resolution spatial sequencing platform (StereoSeq, STOmics), which yielded maps of host and microbiome at 0.5 µm resolution (**Fig. 1a**). This method was performed on tissue sections adjacent to those profiled by Visium. Following the same analysis workflow (**Methods**) as for the low-resolution platform, we mapped host coding and non-coding gene expression, along with microbiome RNA including bacterial rRNA and mRNA. We found that *in situ* polyadenylation again improved the capture of non-coding RNAs and microbes (**Fig. S5**). We confirmed that the measurements performed at low resolution (Visium) and high resolution (StereoSeq) for both host and microbial RNA were in good agreement at the bulk level (**Fig. S6**).

In a section of mouse ileum, we recovered a total of 4.8 million host RNAs (3.77 host UMIs per µm^2^ in the tissue) representing 28,391 genes, and 10 million microbial RNAs (9.19 molecules per µm^2^ in the lumen) representing 81 species with >0.01% abundance (**Fig. S7**). To create a detailed map at the cell level of the host tissue, we used paired imaging data to assign host RNAs to individual cells. We then predicted cell types via computational deconvolution using single-cell RNA-seq data from the same mouse model as a reference^23,24^. Finally, we combined the host map with the microbial signal at 0.5 µm resolution to generate a highly detailed view of the host-microbiome interface (**Fig. 3a**).

**Figure 3.**
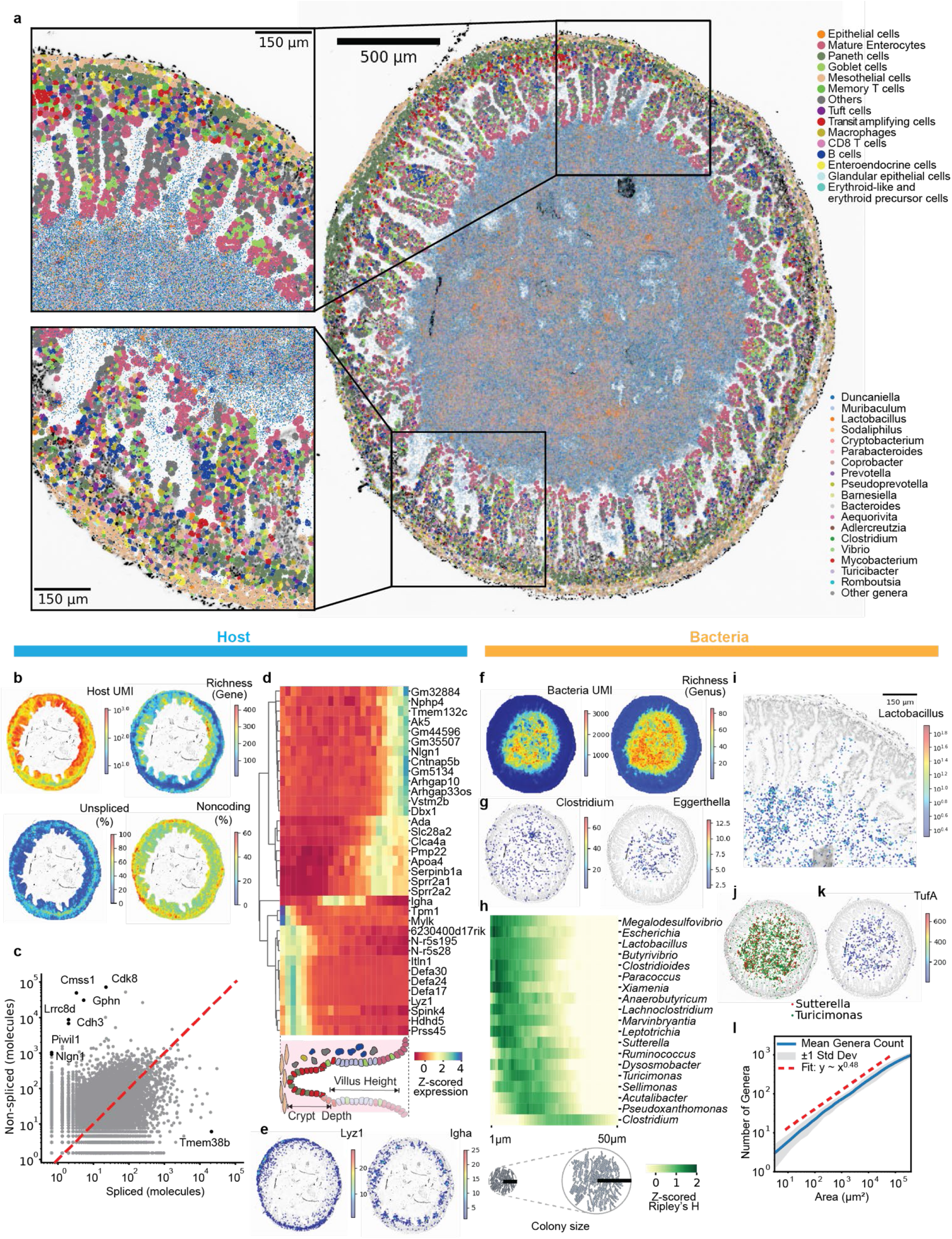
High-resolution spatial mapping of host total gene expression and the microbiome. **a**. Spatial maps of host gene expression and microbiome composition (spots with more than 1 microbial RNA detected are shown). **a**. Spatial maps of host UMIs, gene richness, unspliced RNA and non-coding gene ratio (20 µm square bins), **c**. Plots of spliced and unspliced molecules for each coding gene (outliers in black, points with distance from y=x greater than five standard deviations). **d**. Heatmap of the expression of select genes along the distance from the outer tissue edge. **e**. Spatial gene expression of select genes (20 µm^2^ bins). **f**. Maps of measured unique bacterial molecules (UMI) and genus richness (20 µm^2^ bins). **g**. Spatial maps of abundance of specific genera (20 µm^2^ bins). Spots with more than 1 microbial count are shown. **h**. Z-scored Ripley’s H score. **i**. Zoom-in of abundance of Lactobacillus. **j**. Example of spatially correlated genera. **k**. Spatial maps of bacterial functional gene. **l**. Species accumulation curve for bacterial genera in the ecosystem of the gut.

We observed distinct zonation patterns of coding and non-coding host gene expression. Host gene expression was spatially non-uniform, with significantly higher levels and diversity of gene expression observed at the tips of the villi, likely due to the increased transcriptional activity of mature enterocytes (**Fig. 3b, Fig. S8**). Unspliced mRNAs accounted for 22.5% of the total host RNA in the tissue, with a higher proportion observed at the bases of the crypts, possibly associated with the turnover of transit amplifying (TA) cells (**Fig. 3a**). Genes with a high proportion of unspliced molecules included Cdk8 (Cyclin-dependent kinase 8), a transcription regulatory protein and oncogene associated with human colorectal cancer (**Fig 3c**). These observations are in line with previous studies which have shown that Apc-deficient CRC cells dysregulate RNA splicing machinery^25^. *In situ* polyadenylation improved the capture of non-coding genes in comparison to conventional STomics protocol (**Fig. S5c**). Non-coding RNA expression was elevated in the zone closer to the gut wall. Some non-coding genes showed cell-type specific expression, such as *Hnf1aos1* (**Fig. S9**). We detected several landmark genes, including the non-coding gene *6230400D17RiK*, which was enriched closer to the gut wall, and *Ada*, which was found at the tips of the villi (**Fig. 3d**). Finally, we identified transcripts of genes that are known to be involved in host response to the microbiome, such as the lysozyme encoding gene *Lyz1* and other antimicrobial peptides including defensins expressed by Paneth cells at the base of crypts, as well as Igha, which encodes a segment of the IgA heavy chain, expressed by plasma blasts in the lamina propria (**Fig. 3e**).

The density and diversity of bacteria captured micro-scale ecological features of the lumen. We found that bacterial RNA transcripts were non-uniformly distributed inside the lumen (**Fig. 3f**). We identified fewer bacteria near the boundary with the host, and the bacterial diversity measured at the genus level was also lower close to the boundary with the host. *Clostridium* was evenly distributed in the tract with the exception of one large cluster in the lumen. *Klebsiella* was abundant near the tip of the villi, and *Eggerthella* was abundant away from the host tissue (**Fig. 3g**). We observed colony-like local accumulations for several genera: 54 genera showed significant autocorrelation (moran’s I p-values < 0.05, major genera with >0.01% total bacterial counts) in line with colony-formation (**Fig. S10**). For these genera, we calculated Ripley’s H to infer cluster size (**Fig. 3h**). Some genera including *Lactobacillus* showed small colony size (radius < 10 µm, **Fig. 3i**), while other genera including *Turicimonas* had medium-sizes colonies (∼ 10 µm), and taxa including *Clostridium* formed bigger colonies (> 30 µm). Analysis of spatial correlation between colony-forming genera revealed strong correlations between bacterial genera, including between *Turicimonas* and *Sutterella* (**Fig. 3j)**. The size of bacterial colonies may be influenced by factors such as bacterial reproductive capacity, the abundance of available resources, and the level of intertaxa competition. Further investigation is needed to elucidate how these colonies form over a few hours of passage through the small intestine and how they contribute to microbial community structure.

We aligned bacterial reads to the full rRNA operon database^26^, and estimated that 49.1 % of bacterial reads are non-ribosomal. We annotated these non-ribosomal reads to bacterial genes predicted from assembled bulk metagenomic data measured on sister sections (Methods). 3.7% of these reads were annotated to metagenomic genes including TufA gene (Translation elongation factor EF-Tu, a GTPase, **Fig. 3k**). EF-Tu catalyzes the binding of aminoacyl-tRNAs to the ribosome during translation. It is one of the most abundant and highly conserved bacterial proteins, and indeed was observed to be widely expressed within the lumen.

We measured the relationship between habitat area size and the number of unique species identified in the ecosystem of the mouse gut (**Fig. 3l**). The relationship between the number of unique genera observed and the area sampled followed a power law over three orders of magnitude (16 µm^2^ - 0.16 mm^2^), in line with observations of species-area relationships in a wide range of systems, including plant and animal ecosystems. The observed power exponent of 0.48 (genus level) indicated relatively high spatial dispersion for the microbiome in the ileum of mice relative to exponents reported for plant, animal and environmental microbial ecosystems^27^.

Microbes are an inherent part of the microenvironment of cancers that develop at epithelial barrier surfaces^28^. To study the spatial organization of host cells and gut microbes associated with tumors, we assayed a section of ileum tissue with notable tumors. We compared the microbiome at the edges of tumor and normal tissue (**Fig. 4a, Fig. S11**). To this end, we first defined the boundary between host and microbiome based on microscopy images (**Fig. S12)**, and then measured the spatial organization of host cell types and key taxa as a function of distance to this boundary. This analysis showed that in normal tissue, microbes are most dense 100-200 µm from the host villi (**Fig. 4b**, top), whereas in the tumor tissue, microbes are most dense directly at the boundary with the tumor (**Fig. 4b**, bottom). *Clostridium*, the most abundant genus, and *Lactobacillus* and *Parabacteroides*, were closely associated with the tumor edge (**Fig. 4c**), whereas for normal tissue, these taxa were found away from the tissue boundary towards the lumen. Both *Lactobacillus* and *Turicimonas* again showed evidence of colony formation (radius 10-20 μm, **Fig. S13**). In normal tissue, mature enterocytes were located closest to the host-microbe boundary, followed by immature enterocytes and other intestinal epithelial cells. Paneth cells were located at the basal region, in line with the known architecture of ileum tissue. In contrast, tumor-associated TA cells, and immune cells including dendritic cells, macrophages, and CD8 T cells were enriched in the tumor (**Fig. 4d, Fig. S14a**). At the gene level, expression of some genes moved more toward the lumen in the tumor subregion, including cancer-related genes such as FMNL2 (**Fig. S14b**). Mucin-producing goblet cells were located away from the tissue-microbiome boundary due to the presence of the tumor mass, and consequently, the protective barrier of mucin may not be functioning on the tumor surface. This change in host architecture likely explains the dramatic change in local microbiome composition along the edge of the tumor. Collectively, these data and analysis demonstrate the possibility to map microbe-microbe and microbe-host interactions at high resolution using *in situ* polyadenylation combined with spatial RNA sequencing.

**Figure 4.**
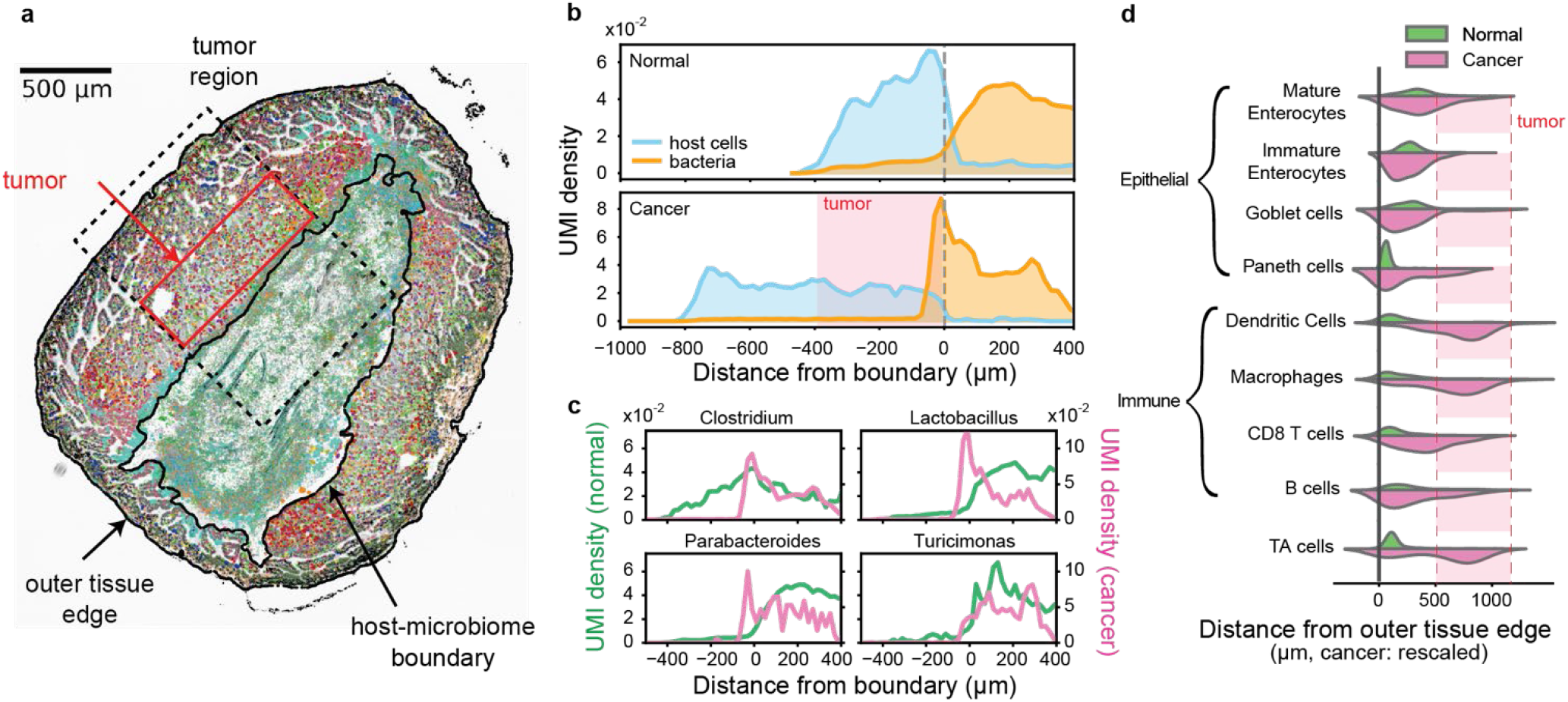
High-resolution spatial mapping of a tumor-microbiome interface. **a**. Spatial map of host gene expression and microbiome in an ileum section with tumor (ApcMin/+ mouse, spots with more than 1 microbial count shown). Color legend is the same as in Figure 3a. **b**. Probability density plot of host and microbial cells as function of distance to the host-microbiome boundary, for normal (top) and tumor laden (bottom) tissue. **c**. Genera abundance as function of distance to the host-microbiome boundary for normal (green) and tumor laden (pink) tissue. **d**. Violin plot of cell density as function of distance from the outer tissue edge. The distances of each cell type in the tumor were linearly rescaled to match the location of mature enterocytes in a section without tumor.

## DISCUSSION

Characterizing the spatial organization of microbes in the gut is crucial for understanding the inter-microbial and host-microbial interactions that govern the organ-like function of the gut microbiome. Yet, current methods for mapping the gut microbiome have significant limitations. In this study, we show that combining *in situ* polyadenylation with spatial RNA-seq effectively maps the biogeography of the gut microbiome and the host A-tailed and non-A-tailed transcriptomes. By integrating a simple enzymatic step with commercially available spatial transcriptomics platforms, this method provides an accessible and scalable way to measure the host-microbe interactome across spatial scales.

We applied this methodology to profile intestinal tissue in a mouse model of intestinal neoplasia. We first demonstrated this method at low spatial resolution and characterized changes in microbiome composition as a function of longitudinal and transverse location in the mouse intestine, which corroborated many previously known features of the organization of the gut microbiome in mice. The enhanced recovery of microbial RNA enabled by *in situ* polyadenylation then allowed high-resolution, 0.5 μm spatial sampling of the microbiome and host total RNA expression. This high-resolution analysis revealed interactions within and between microbial taxa by enabling the measurement of spatial heterogeneity and colony formation, even for colonies less than 10 μm in radius. Colony formation may indicate active growth, and if so, colony size may be a proxy for growth rate, especially in the colon where mixing is reduced. We also observed mechanisms by which the host tissue architecture changes local microbiome composition. At the boundary between the microbiome and tumors, we observed a pronounced shift of key microbial taxa towards the boundary with the host, suggesting increased (generalized) host-microbe interactions. These changes in local microbiome structure are likely explained by the altered local host architecture, with mucin-producing cells dislocated from the tissue boundary.

Importantly, as we have shown previously, *in situ* polyadenylation enabled mapping of both the A-tailed and non-A-tailed host transcriptome. Analysis of the “total” transcriptome revealed spatially restricted expression of several classes of noncoding RNAs, reflecting the architecture of intestinal tissue. We identified landmark coding and non-coding molecules along the crypt-villus axis, with increased expression in mature enterocytes at the villus lining and a higher fraction of unspliced, newly transcribed RNA near the crypt. These patterns point to the potential of using this assay to study intestinal stem cell differentiation dynamics.

While this study lays the groundwork for consideration of spatial structure in microbiome research, there are limitations that need to be addressed. First, the high cost of commercial spatial transcriptomics platforms remains significant, hindering broader adoption and use of these techniques in drug screening applications. Second, long-read sequencing could enhance taxonomic classification beyond what is possible with short-read sequencing alone, and may enable further analyses, for example spatial profiling of the gut immune repertoire. Last, making the methodology compatible with formalin-fixed paraffin-embedded tissue would open application of these techniques in pathology^29^. Despite these limitations, this study shows that spatial transcriptomics provides a unique window into microbiome ecology and intermicrobial and host-microbial interaction. Going forward, spatial transcriptomics will be a powerful approach to explore questions in gut immunology^30,31^, to explore microbial colonization of mucus and intestinal tissue, to study microbiomes in small niches such as crypts, and to investigate the concept of the cancer-associated microbiome. Spatial transcriptomics can further be applied to explore the role of specific taxa in diseases with known microbiome involvement, such as inflammatory bowel disease and other autoimmune disorders. Ultimately, spatial transcriptomics addresses an unmet need by enabling simultaneous *in situ* profiling of both host and microbiome at high resolution, allowing for the survey of structural relationships from the macroscale to the microscale.

## METHODS

### Animal Models and Experimental Procedures

All animal protocols were approved by the Cornell University Institutional Animal Care and Use Committee (IACUC), and experiments were performed in compliance with institutional guidelines. C57BL/6-ApcMin/+/J mice were used for the spatial transcriptomics experiments. All mice (C57BL/6-ApcMin/+/J and C57BL/6-Wild type) were maintained at the barrier mouse facility at Weill Hall of Cornell University. ApcMin/+ and wild-type mice were initially ordered from Jackson Laboratory and then bred in the barrier facility. The ApcMin/+ mice used in these experiments have a chemically induced transversion point mutation at nucleotide 2549, resulting in a stop codon at codon 850, truncating the APC protein.

Both male and female mice were used, and their precise age was noted. Experimental and breeding mice were provided with *ad libitum* access to autoclaved water and rodent chow (autoclavable Teklad global 14% protein rodent maintenance diet #2014-S; Envigo). The overall health, food intake, and weight of the mice were closely monitored to ensure that tumor burden did not violate ethical standards. After approximately 100 days, the mice were sacrificed using 5 minutes of CO2 asphyxiation followed by tissue collection. The intestines from the mice were inspected for tumor localization, and excess fat was removed. The intestines were then cut into individual sections, embedded in cryomolds with O.C.T Compound (Tissue-Tek), and frozen in an isopentane-liquid nitrogen as described previously<u>^11^.</u> Specifically, the small intestine was cut into 4-6 approximately equal-sized segments, the large intestine into 2-3 segments, and the cecum was processed separately.

### In situ polyadenylation for the gastrointestinal tract profiling with the Visium platform

Cryosections were obtained from four distinct locations of the intestine of the same individual (male, 13w) — the proximal small intestine, ileum, cecum, and large intestine. Sections were processed using either a modified protocol or the standard Visium protocol. For the modified protocol, 10 µm thick tissue sections were mounted onto Visium Spatial Gene Expression v1 slides. The sections were fixed in freshly prepared methacarn solution (60% methanol, 30% glacial acetic acid, 10% chloroform) at room temperature for 15 minutes. H&E staining was performed according to the Visium protocol, and tissue sections were imaged using a Zeiss Axio Observer Z1 microscope equipped with a Zeiss Axiocam 305 color camera. The resulting H&E images were corrected for shading, stitched, rotated, thresholded, and exported as TIFF files using Zen 3.1 software (Blue edition). After imaging, the slides were transferred into the Visium Slide Cassette.

*In situ* polyadenylation was conducted using yeast Poly(A) Polymerase (yPAP; Thermo Scientific, Cat #74225Z25KU). Each capture area was equilibrated by adding 100 µl of 1X yPAP Reaction buffer (20 µl 5X yPAP Reaction Buffer, 2 µl 40U/µl Protector RNase Inhibitor, 78 µl nuclease-free H_2_O), incubating at room temperature for 30 seconds, and then removing the buffer. Following this, 75 µl of yPAP enzyme mix (15 µl 5X yPAP Reaction Buffer, 3 µl 600U/µl yPAP enzyme, 1.5 µl 25 mM ATP, 5 µl Murine RNase Inhibitor, 50.5 µl nuclease-free H_2_O) was added to each reaction chamber. The chambers were sealed and incubated at 37°C for 25 minutes, after which the enzyme mix was removed. Post-polyadenylation, a 30-minute enzymatic permeabilization step was performed, followed by the standard Visium library preparation protocol to generate cDNA and final sequencing libraries. For the standard Visium experiment, H&E staining and imaging were immediately followed by permeabilization and the standard library preparation.

### In situ polyadenylation with the STOmics platform

Adjacent ileal cross-sections to those profiled with Visium were also profiled using either the modified or standard STomics protocol. 10 µm thick sections were placed onto STOmics mini chips (Product No. 211ST004). For the modified protocol, sections were fixed in methacarn for 15 minutes as previously described, followed by a DNA staining step according to the STOmics protocol. Imaging was performed on a Zeiss Axio Observer Z1 Microscope using a Hamamatsu ORCA Fusion Gen III Scientific CMOS camera. Images were stitched, rotated, thresholded, processed, and exported as TIFF files using Zen v.3.1 software (Blue edition), and then registered using the STOmics software. After imaging, *in situ* polyadenylation was performed followed by 12 minute permeabilization and library preparation according to the STOmics protocol. For the standard experiment, imaging is directly followed by permeabilization. Additionally, ileal cross-sections from a second mouse (female, 17w), containing a tumor adjacent to the luminal cavity, were processed exclusively using the modified protocol.

### Sequencing of the spatial transcriptomics libraries

Sequencing of the Visium libraries was performed on a NextSeq 2K (Illumina) platform using a P3 200bp kit, with reads allocated as follows: 28 bp for read 1, 10 bp for index 1, 10 bp for index 2, and 190 bp for read 2. For the libraries prepared using the STOmics platform, sequencing was carried out on a Complete Genomics DNBSEQ-T7 Sequencer using the DNBSEQ-T7 High-throughput Sequencing Set (FCL PE100) and the associated STEROmics primer set. The sequencing run consisted of a 50 bp read 1 (with dark cycles from bases 26 to 40), a 100 bp read 2, and a 10 bp index read.

### Preprocessing and alignment of spatial transcriptomics data

To ensure similar alignment and quantification across platforms and methodologies we used the “slide_snake” pipeline that utilizes Snakemake^32^ (6.1.0), which can be found on github (https://github.com/mckellardw/slide_snake). For the Visium and STRS (Visium) libraries, the pipeline first trims poly(A) and poly(G) sequences, as well as primer sequences using cutadapt^33^. The reads were aligned using STAR v2.7.10a^34^ and STARSolo^35^ (specified parameters: --outFilterMultimapNmax 50, -- soloMultiMappers EM, --clipAdapterType CellRanger4) to generate expression matrices for every sample. For downstream analyses the GeneFull matrices were used. Barcode whitelists and the associated spot spatial locations for Visium data were copied from the Space Ranger software (“Visium-v1_coordinates.txt”). For the StereoSeq and STRS (StereoSeq) libraries, barcode maps were provided by the manufacturer as .h5 files and converted to text format using ST_BarcodeMap (https://github.com/STOmics/ST_BarcodeMap). Alignment references were generated from the GRCm39 reference sequence using GENCODE M32 annotations.

### Unmapped reads classification and construction of microbiome Anndata objects

In this study, to classify reads of microbial origin out of the unmapped reads we utilized Kraken2 (version 2.09)^18^. We used the standard Kraken2 database supplemented with the mouse genome. Unmapped reads flagged in the BAM file were processed to retain the correct cell barcode and unique molecular identifier (UMI) information as identified by STARsolo. This allowed for the demultiplexing of Kraken2 output by cell barcode and UMI. For data integration, we employed Pandas, Scanpy, NumPy, Scipy, and regular expressions to create an AnnData object with cell barcodes as observations and NCBI taxonomy IDs as features. Only classified reads were retained for subsequent analysis.

### Sterile control pre-processing and identification of taxa to filter

To assess the Kraken2 classified microbial counts occurring in non-intestinal tissues for the low-resolution platform we re-aligned previously published Visium and STRS libraries of mock-infected C57BL/6J 11 days year old mice with and without polyadenylation as described in the corresponding studies^11,17^. 85 taxa occurring at 1ppm (UMI) or greater were excluded from downstream analysis as potential misclassification. For the Stereoseq libraries, a sterile control experiment was conducted. Briefly, fresh-frozen heart from a eleven day old mouse were sectioned on a Stereo-seq 1cm × 1cm tile (STOmics, BGI). The sample was fixed in methanol at -20°C for 20 minutes followed by the *in situ* polyadenylation and the STomics library preparation protocol as described above. Taxa occurring at frequencies higher than 1 ppm UMI were excluded from downstream analyses.

### Pre-processing of the Visium and STRS data

Spatial coordinates were assigned to the Visium and STRS library spots based on the barcode map provided by the Space Ranger software (“Visium-v1_coordinates.txt”). The accompanying hematoxylin and eosin histology images of each experiment were used to manually mark the spots that correspond to tissue and lumen. Scanpy^36^, mudata^37,38^, and muon^37^ were used to construct multimodal objects separately for the microbial maps (in the taxonomic levels of phylum, family, genus, and species). This was done for each one of the accounted microbial superkingdoms of Archaea, Bacteria and Viruses. For downstream analyses, only the spots covered by tissue or corresponding to lumen were accounted for.

### Microbial percentage and enrichment calculation for the paired Visium STRS experiments

For the three discussed superkingdoms, the percentage of reads falling under to a superkingdom classification was calculated as the percentage of Kraken-classified reads that belong to the superkingdom over the total counts of the library defined as the sum of unique molecules aligned to the host and unique molecules classified by Kraken2. The enrichment for each paired experiment was defined as the ratio of those percentages.

### Relative abundance and bacterial richness calculations for the low-resolution datasets

To calculate the relative abundance for each examined sample, at family level, the corresponding family reads were collapsed and divided by the total molecules originating from bacteria as classified by Kraken2. The microbial richness per spot was calculated as the number of unique taxa occurring per spot after the exclusion of taxa accounting for 0.01% or less of microbial molecules in the whole sample. For the transverse axis relative abundance analysis, cells were spatially binned from the tissue to the lumen based on their minimum distance to the lumen-associated region. Phyla relative abundance data were then aggregated within each bin to quantify relative abundances across the tissue-lumen axis.

### Cell type deconvolution

We employed the cell2location^24^ model (version 0.1.3) to deconvolve spatial transcriptomics data for the experiments conducted with both Visium and StereoSeq technologies. The scRNA-seq reference, derived from a previous study on Apc Min/+ mice^23^ was filtered to include only genes that are highly expressed and informative for identifying rare cell types, with thresholds set at cell_count_cutoff = 5, cell_percent_cutoff = 0.01, and nonz_mean_cutoff = 1.12. Cell-type-specific expression signatures were generated using negative binomial regression from these selected genes. These signatures were applied to the spatial transcriptomics data to determine cell-type identities, with the highest prediction scores used for assignment. For Visium, we set N_cells_per_location to 30, and for StereoSeq, we set it to 1, with the detection_alpha parameter set to 20 in both cases.

### Bacterial gene function analysis

Considering the low annotation efficiency of the functional composition of the mouse metagenome, we used genes identified from the metagenome data obtained from the same sample as a reference. We predicted the genes from contigs of metagenomic data from the same sample using prodigal^39^ (v2.6.3) and then clustered them with CD-HIT^40^ (v4.6.4) to create a gene reference. The genes in the created gene database were annotated with EGGNOG database (v5.0) using DIAMOND^41^ (v2.0.13) with e-value <1e-5. Meanwhile, in the previous section, reads annotated as Bacteria by Kraken2 were further mapped to the full-length rRNA operon database using BLASTn (identity >80, coverage >60). Unmapped reads were then mapped to the gene database created from the metagenomic data for gene annotation.

### Spatial autocorrelation analysis

Moran’s I was calculated for the major genera (abundance > 0.01%) using the Moran function from the Python library pysal. Spatial weights were generated using the *k*-nearest neighbors (KNN) matrix (*k*=4) from the weights module in pysal. For genera with a Moran’s I p-value < 0.05, Ripley’s H was subsequently derived using the following formula:

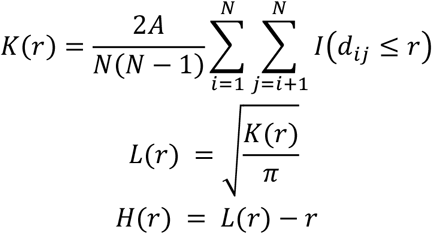

where *d*_*ij*_is the Euclidean distance between points i and j. *I*(*d*_*ij*_ ≤ *r* )is an indicator function that is 1 if the distance *d*_*ij*_ is less than or equal to *r*, and 0 otherwise. *A-*is the area of the observation window. *N-*is the number of points in the dataset.

### Boundary detection

The microscope data was saved in grayscale and then averaged using the OpenCV blur function with a kernel size 100 µm. After that, the data was binarized with a threshold of 80 for normal tissue and 100 for cancer tissue. Finally, boundaries were extracted using the OpenCV findContours function.

## DATA AVAILABILITY

Data will be made available upon publication under GEO accession numbers; GSE276866 for the low-resolution datasets, GSE277196 and GSE277197 for the high-resolution datasets.

## CODE AVAILABILITY

Code associated with this work can be found at https://github.com/ntekasi/microSTRS

## ACKNOWLEDGMENTS

We thank the Cornell Biotechnology Resource Center and Logan Schiller for their help with sequencing the libraries. We thank the Cornell Center for Animal Resources and Education for animal housing and care. We thank Madhav Mantri, Shaowen Jiang, Michael Wang, Rohit Agarwal, and the other members of the De Vlaminck lab for helpful discussions and feedback. We also thank Josh Jones, Shohei Asami, and Wataru Suda for the helpful discussion.

## AUTHOR CONTRIBUTIONS

IN, LT, DWM, and IDV conceived of the study. IN, LT, PS, BMG and QS performed the experiments. IN, LT, DWM, CH and MS analyzed the data. IN, LT and IDV wrote the manuscript. All authors provided input and comments.

## COMPETING INTERESTS STATEMENT

DWM, IN, and IDV have filed a patent on technology described in this work.

## Notes

### Summary of Updates

We have updated the main text to enhance clarity and focus.

