## Supplementary information for "High Resolution Spatial Mapping of Microbiome-Host Interactions via *in situ* Polyadenylation and Spatial RNA Sequencing"

### SUPPLEMENTAL FIGURES

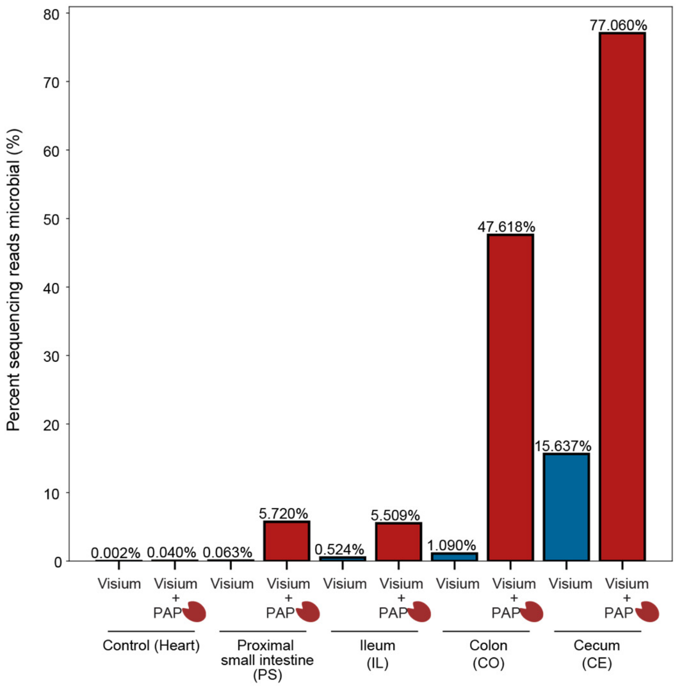


**Supplementary Figure 1.** Percentage of sequencing reads classified as Microbial. The bar plots show the percentage of reads that did not map to the mouse genome and were classified as microbial (belonging to Bacteria, Archaea, or Viruses superkingdoms) for the non-intestine control/ Murine Heart samples Visium and Visium with *in situ* Polyadenylation, and the paired experiments for the 4 examined sites of the gastrointestinal tract. The samples with *in situ* polyadenylation are marked with a red enzyme cartoon.

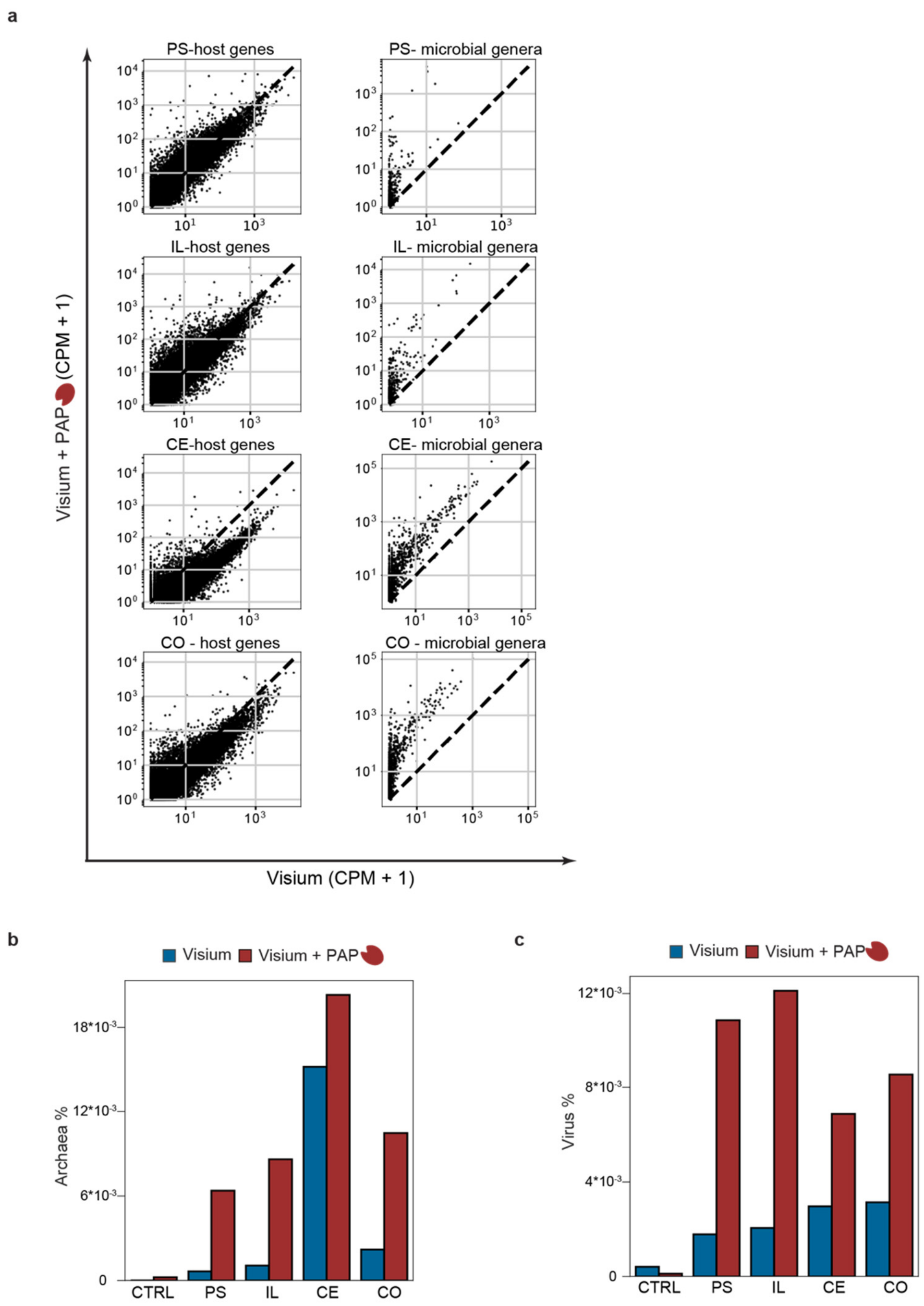


**Supplementary Figure 2.** Enrichment of microbial signal capture via *in situ* polyadenylation. (a) Scatter plots showing the Aggregated unique molecules per million for host genes (left) and bacterial taxa (right) for the tissue pairs. (b)Percent of Unique molecules captured classified as archaea for the paired experiments with (red) and without (blue) *in situ* polyadenylation for the Murine heart and the 4 sampling tissue sites. (c) Percent of Unique molecules captured classified by Kraken2 as viruses for the paired experiments with (red) and without (blue) *in situ* polyadenylation for the Murine heart and the 4 sampling tissue sites.

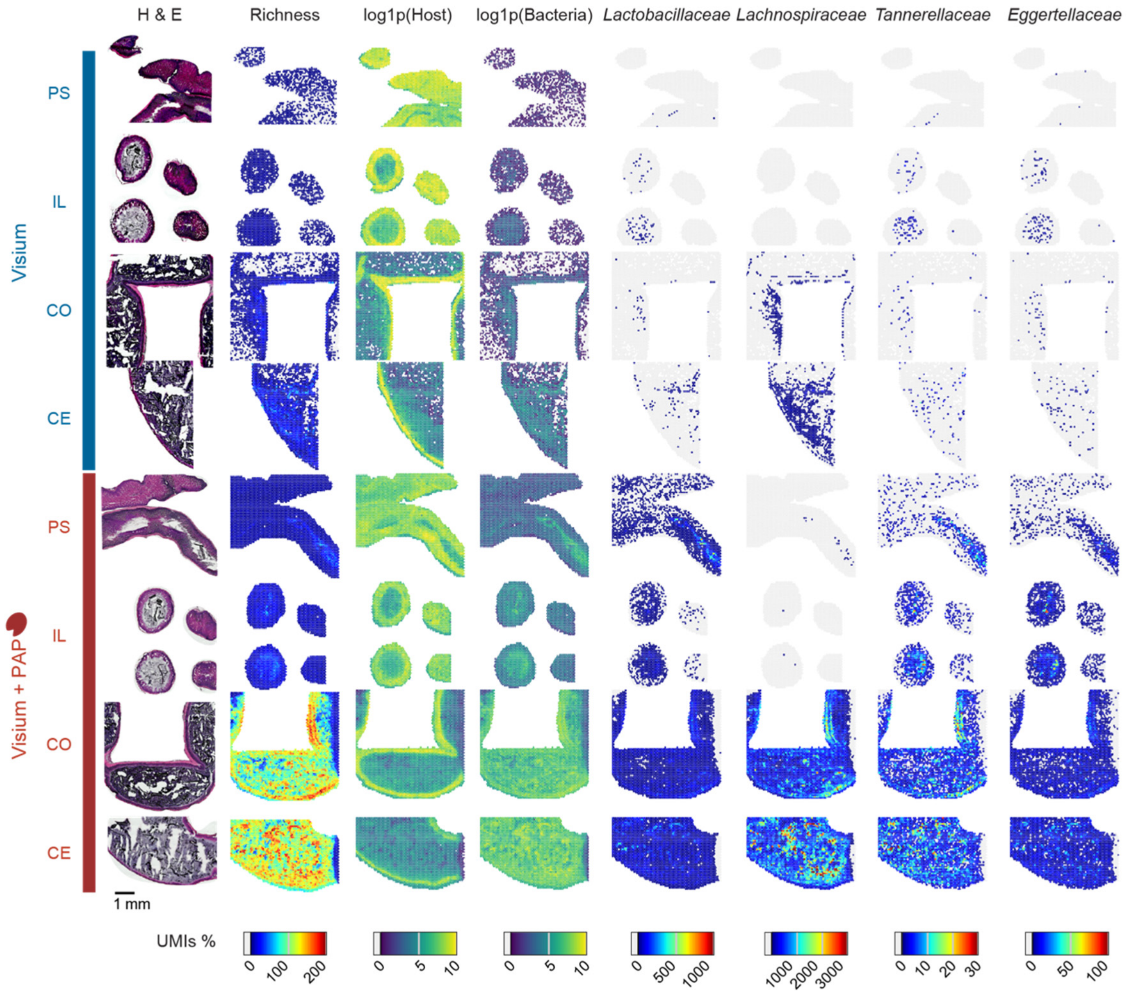


**Supplementary Figure 3.** Spatial feature plots and histological data for the tissues processed with the low-resolution platform (with and without polyadenylation).

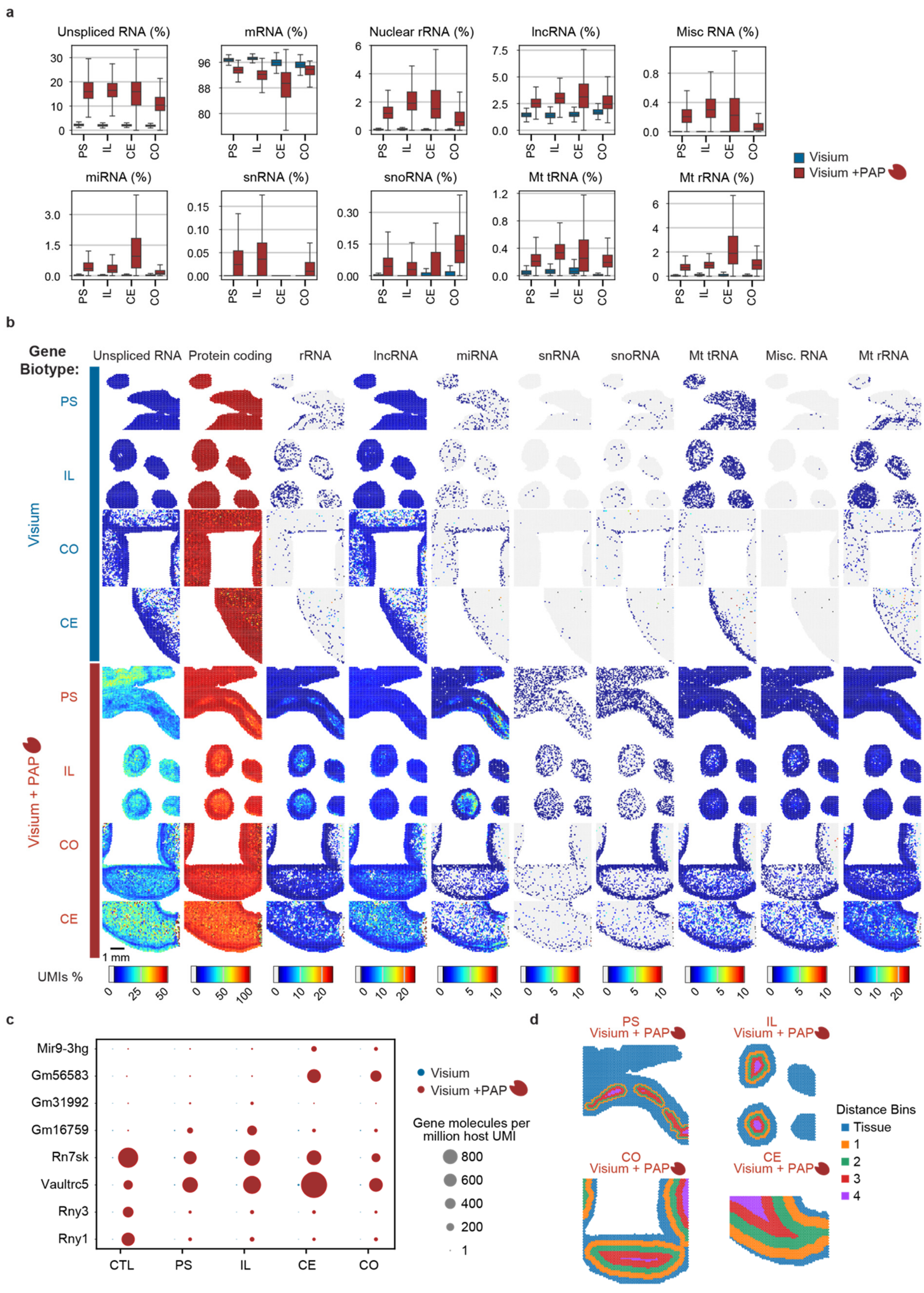
a
Unspliced RNA (%)
mRNA (%)
Nuclear rRNA (%)
lncRNA (%)
Misc RNA (%)
miRNA (%)
snRNA (%)
snoRNA (%)
Mt tRNA (%)
Mt rRNA (%)
PS
IL
CE
CO
Visium
Visium +PAP
b
Gene Biotype:
Unspliced RNA
Protein coding
rRNA
lncRNA
miRNA
snRNA
snoRNA
Mt tRNA
Misc. RNA
Mt rRNA
Visium
Visium + PAP
1 mm
UMIs %
c
Mir9-3hg
Gm56583
Gm31992
Gm16759
Rn7sk
Vaultrc5
Rny3
Rny1
CTL
Gene molecules per million host UMI
800
600
400
200
1
d
PS Visium + PAP
IL Visium + PAP
CO Visium + PAP
CE Visium + PAP
Distance Bins
Tissue
1
2
3
4

**Supplementary Figure 4.** RNA biotype distributions for the paired Visium and STRS experiments. **(a)**Boxplots representing the distributions of select RNA biotypes across the 4 sampling sites. **(b)** Spatial maps showing the patterns of the biotypes across the paired experiments. **(c)** Dotplot of selected genes that are common to all four GI tract regions (PS, IL, CE,CO) and murine heart tissue (CTL). **(d)** Maps showing the spatial bins used for the relative abundance calculation.

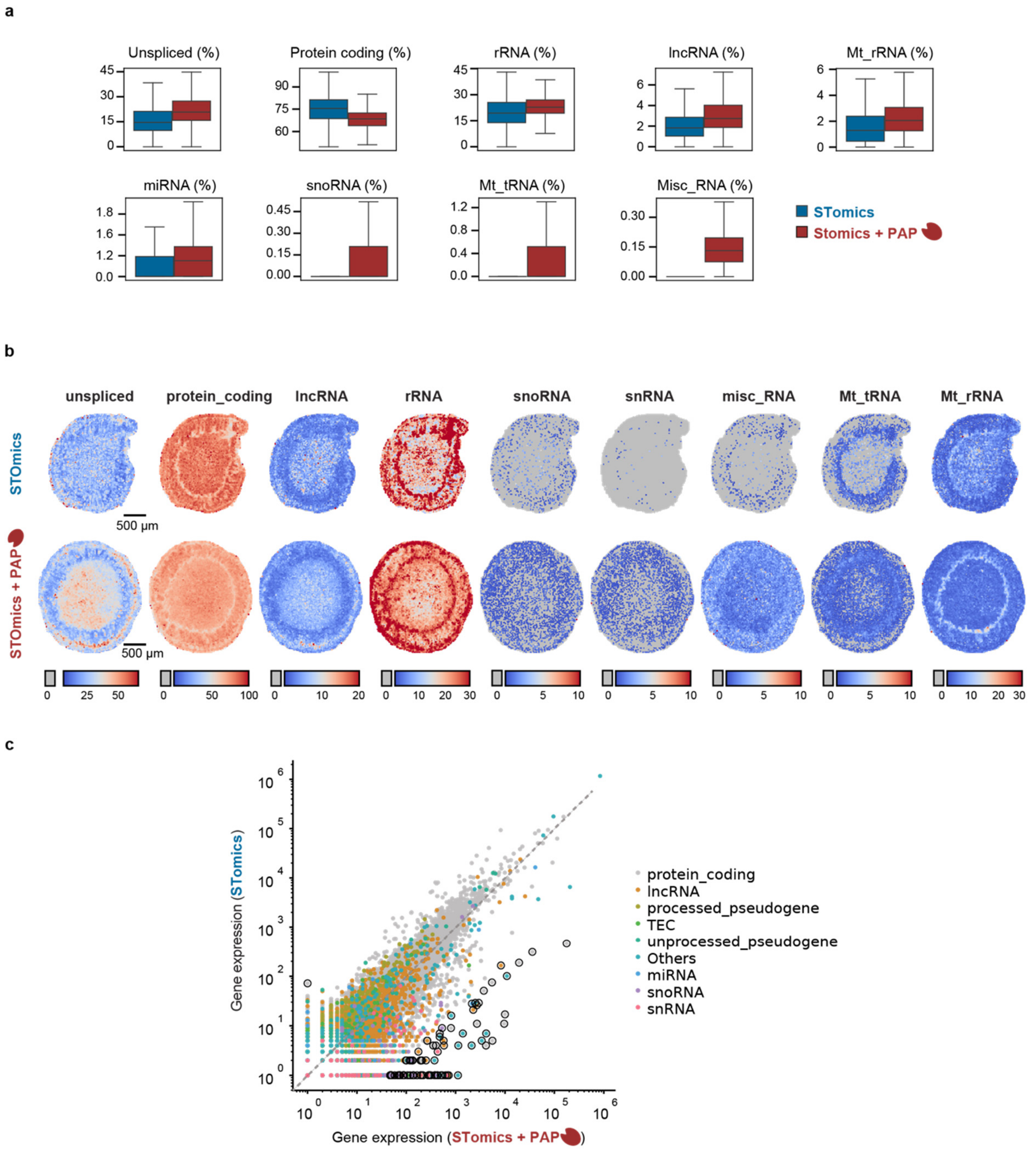


**Supplementary Figure 5.** RNA biotype distributions for the paired Stereoseq and STRS (Stereoseq + PAP) experiments. **(a)** Boxplots representing the amount of select RNA biotypes **(b)** Spatial maps showing the patterns of the biotypes across the paired experiments. **(c)** Scatter plot showing the rescaled sum of unique molecules per million for host genes. The gene expressions were linearly rescaled so that the sum of each expression matched the mean of the sums.

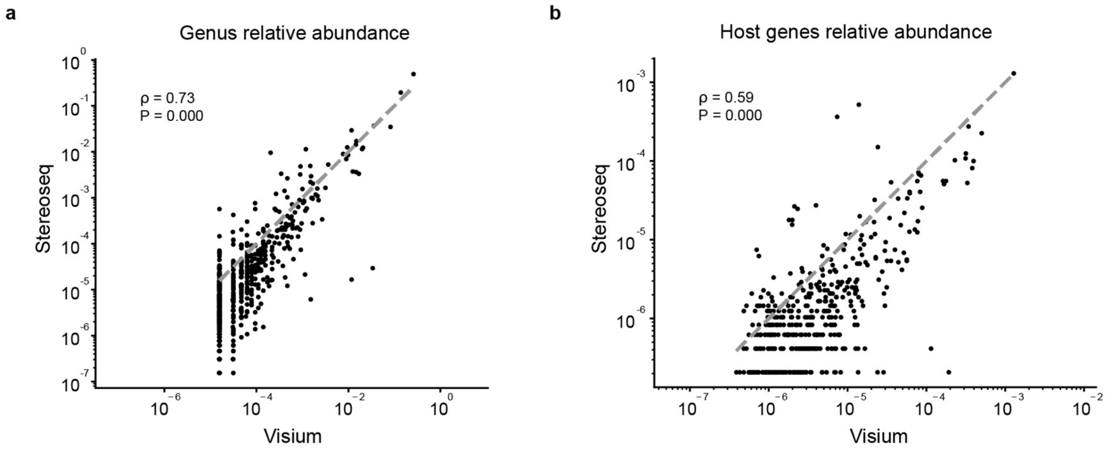


**Supplementary Figure 6.** Comparison between low resolution platform Visium and high-resolution platform Stereoseq. **(a)** Genus-level relative abundance measured on the Visium (*in situ* polyadenylation) and high resolution Stereoseq platform (*in situ* polyadenylation). **(b)** Host gene relative abundance measured on the Visium (*in situ* polyadenylation) and high resolution Stereoseq platform (*in situ* polyadenylation).

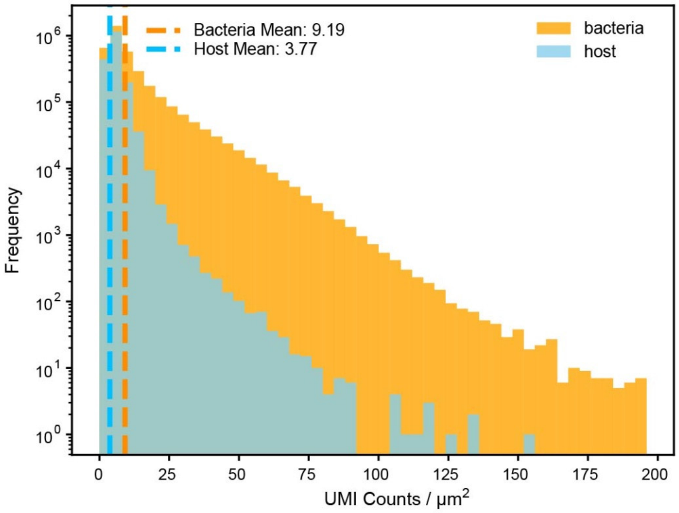


**Supplementary Figure 7.** The frequency distribution and mean of UMI counts per $\mu m^2$ across the whole tissue and lumen areas.

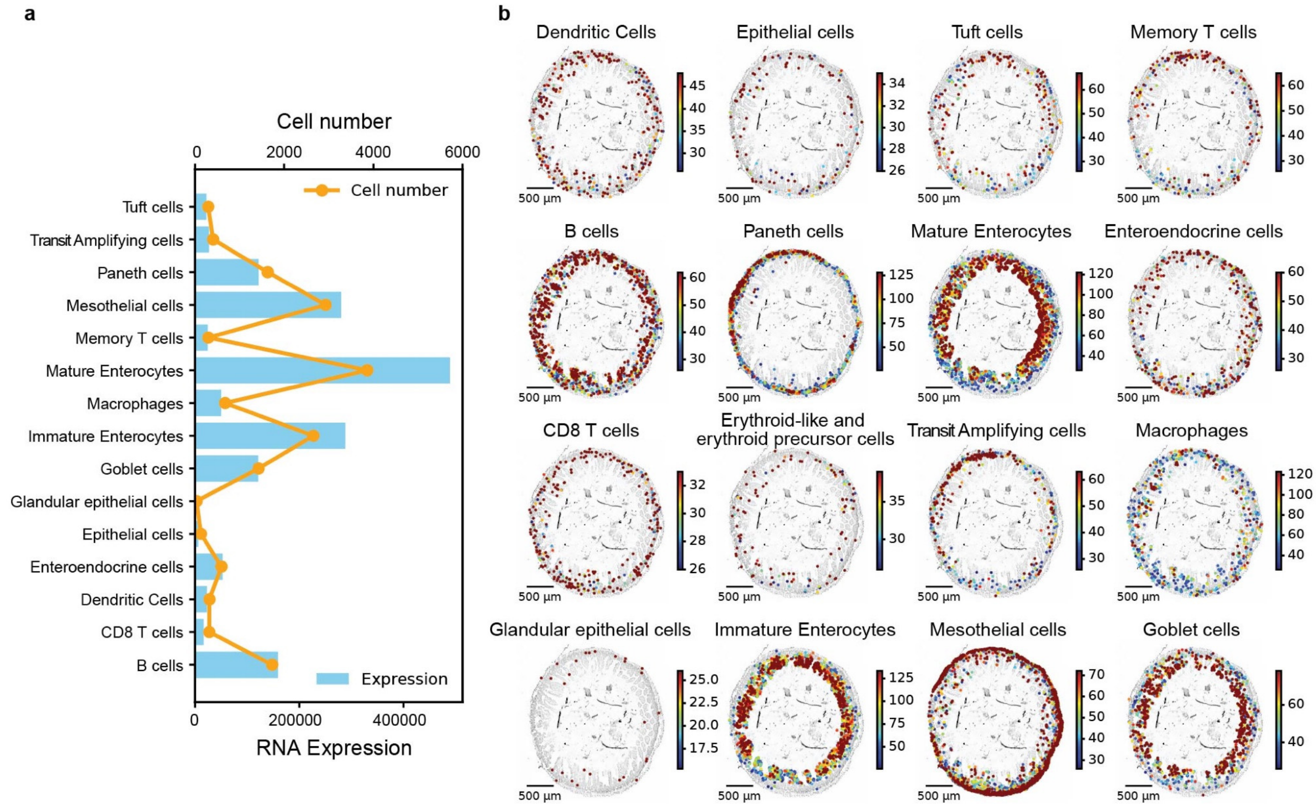


**Supplementary Figure 8.** Cell type analysis. **(a)** The total number of cells and RNA molecules for each cell type. **(b)** Spatial maps of each cell type.

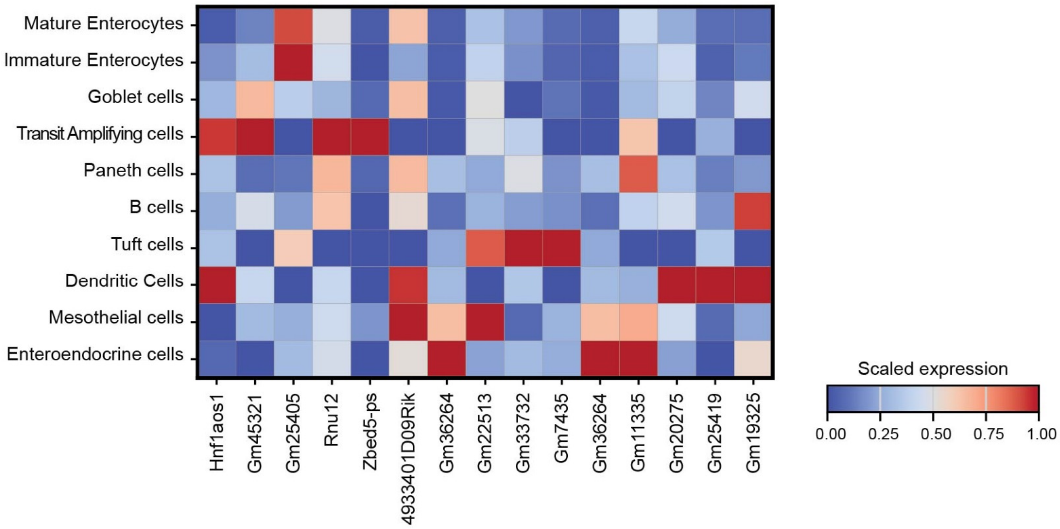


**Supplementary Figure 9.** Heatmap of the normalized amount of non-coding genes for each cell type. We used min-max normalization to ensure that the smallest value for each gene was set to 0 and the largest value was set to 1.

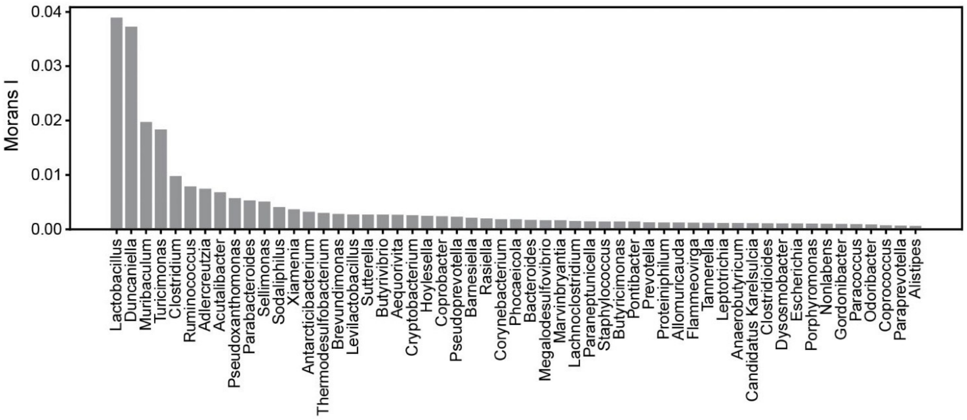


**Supplementary Figure 10.** Spatial autocorrelation and correlation analysis in genera. Moran's I scores of major genera ( >0.01% total bacterial counts, *P*-value < 0.05).

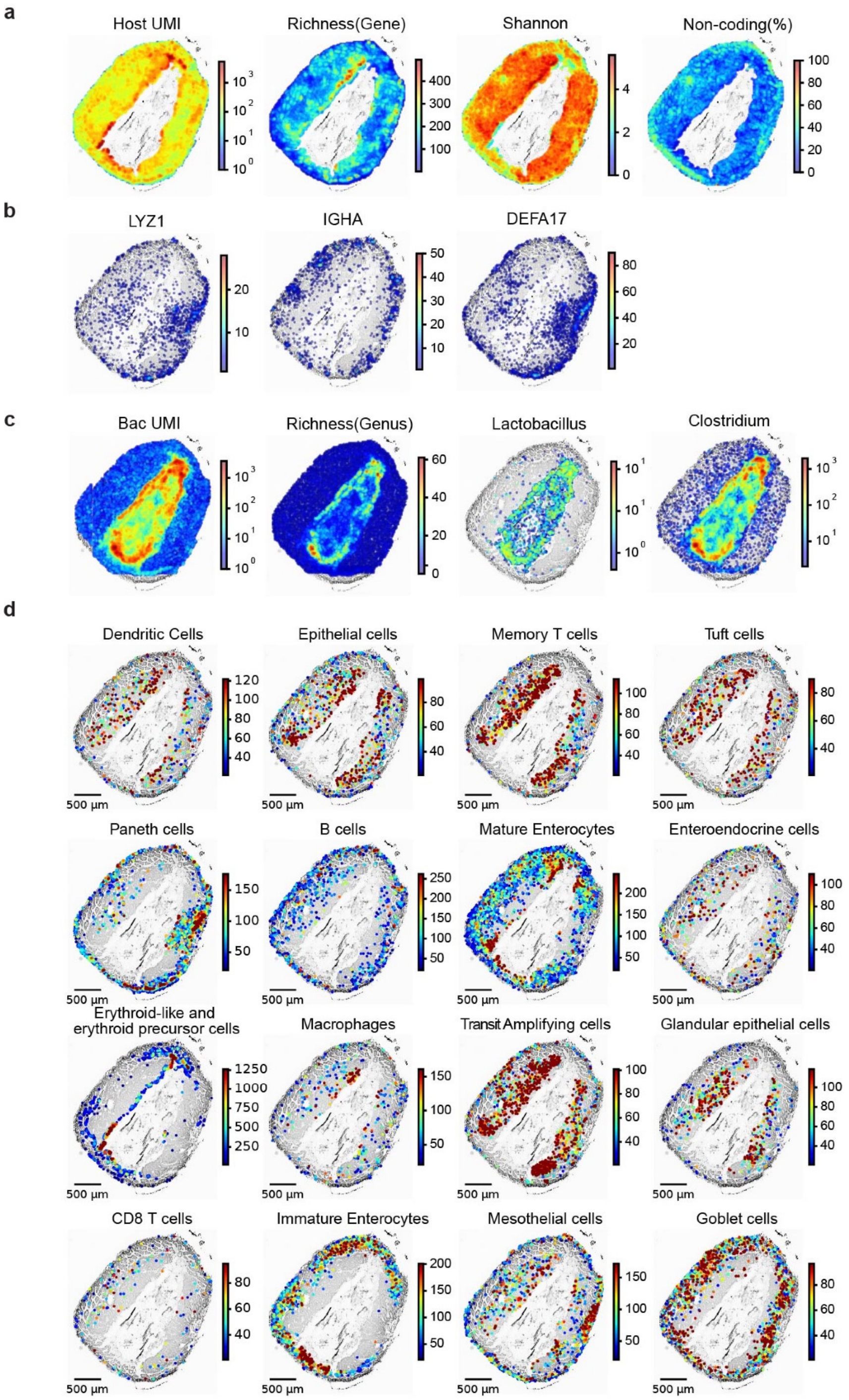


**Supplementary Figure 11.** Spatial mapping of cancer section. **(a)** Spatial mapping of host UMIs, gene richness, unspliced molecule and non-coding gene ratio in 20 µm square bin, **(b)** Spatial mapping of ratio of selected gene expressions in 20 µm square bin. **(c)** Spatial mapping of bacterial UMIs, gene richness in 20 µm square bin. **(d)** The spatial mapping of each cell type.

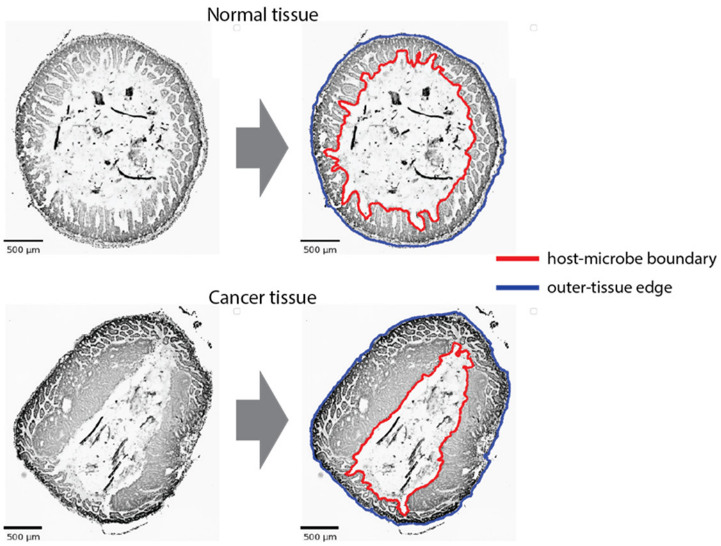


**Supplementary Figure 12.** Defined boundaries for lumen, tissue area based on the microscope images. After average smoothing in 100um, image data were binarized to detect the boundaries for both host-microbe boundaries and out-tissue edges in cancer and normal samples.

**a**

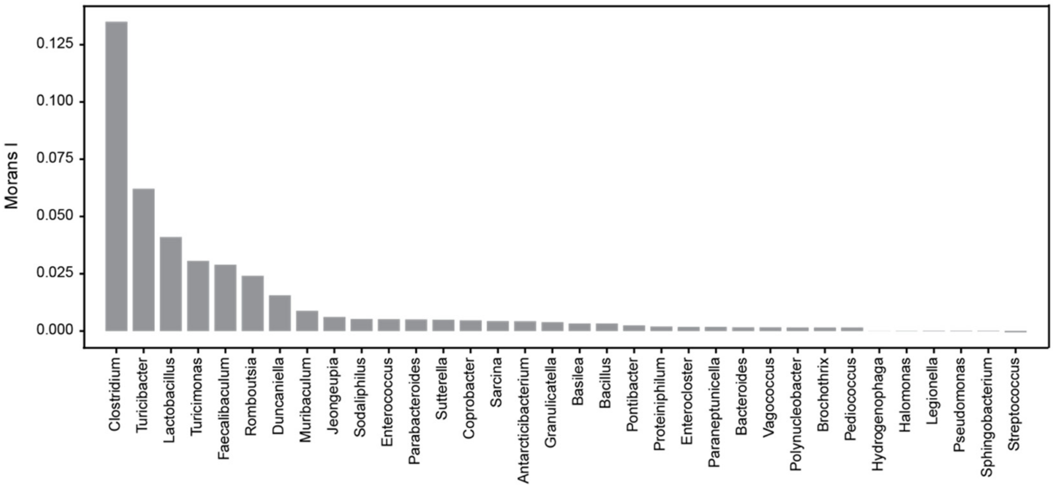


**b**

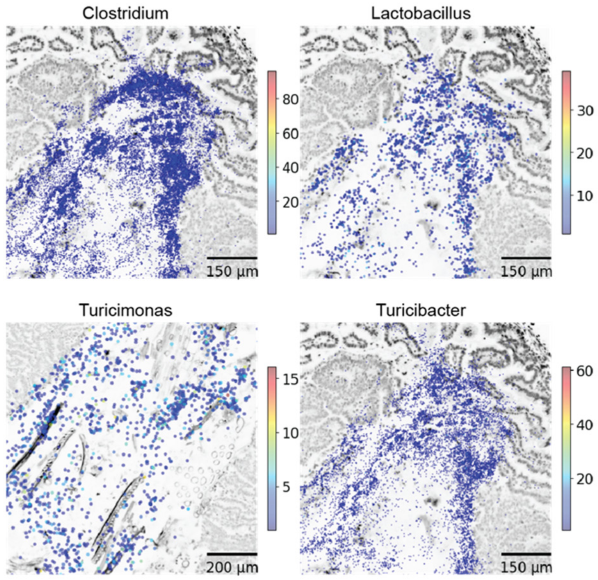


**Supplementary Figure 13.** Autocorrelation in cancer section. **(a)** Moran's I scores of major genera ( >0.01% total bacterial counts, *P*-value < 0.05). **(b)** Enlarged spatial mapping image of selected genera.

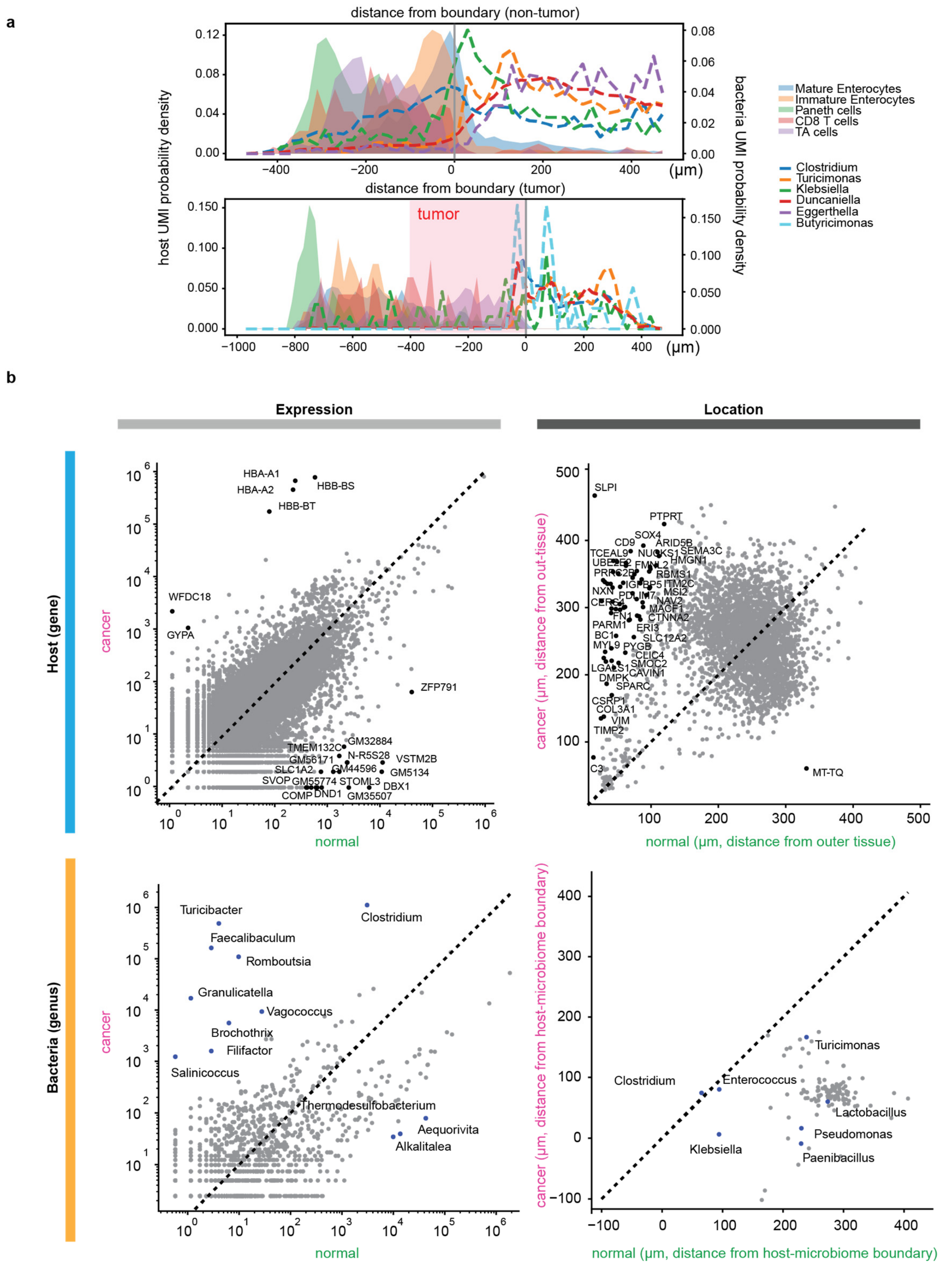


**Supplementary Figure 14.** Difference in location of the host genes and microbes between normal and cancer sections. **(a)** Probability density selected host cells and microbes in the function of the distance from host-microbe boundary. **(b)** Gene abundance of normal and cancer sections(left-above), and genus level abundance in normal and cancer sections (left-below). Median distance of genes from out-tissue area (right-above), and genera from host-microbe boundary (right-below).
